# Biological and molecular characterization of six Shiga toxin-producing *Escherichia coli* (STEC) strains encoding the stx_2_ gene in Colombia

**DOI:** 10.1101/2020.07.17.209239

**Authors:** Brayan Stiven Arango-Gil, Sebastián Peña-Buitrago, Jhon Carlos Castaño-Osorio, Claudia Viviana Granobles-Velandia

## Abstract

Shiga toxin-producing *Escherichia coli* (STEC) is a bacterial pathogen that cause diarrhea and severe human diseases. Its principal virulence factor are the Shiga toxins Stx1 and Stx2 which have been identified diverse subtypes considered to be responsible for severe complications of STEC infection. These toxins are encoded in temperate bacteriophages and their expression is linked to phage lithic cycle, which is regulated by late genes and the Q anti-terminator protein. The aim of this study was to characterize biologically and molecularly STEC strains encoding stx_2_ gene isolated from cattle feces in Colombia. We selected six STEC strains, which were evaluated its Stx production, the Stx_2_ subtypes, induction of the lithic cycle of bacteriophages and its late region. The results evidenced two highlighted strains with high levels of Stx production and induction of the lithic cycle, compared with the others. Likewise, the strains evaluated showed three Stx_2_ subtypes: Stx_2_a, Stx_2_c, and Stx_2_d. Regarding the late region, most of the strains carried the *qO111* allele and only one strain showed differences in the *ninG* gene. Although the sample was limited, variability was observed in the Stx production assay, induction of the lithic cycle, Stx_2_ subtypes and late region of the phages, which could indicate the diversity of the phages carrying STEC strains in Colombia.

## INTRODUCTION

Shiga toxin-producing *Escherichia coli* (STEC), is an emerging pathogen involved in food-borne infections which causes diarrheal disease outbreaks, like hemorrhagic colitis (HC) and hemolytic uremic syndrome (HUS). The HUS is one of the most severe human disease caused for STEC, characterized for producing thrombocytopenia and acute kidney failure, which mainly affect children under 5 years old and older adults [1]. Cattle have been considered the main reservoir of pathogenic STEC for humans and its transmission occurs through the consumption of contaminated foods with feces from cattle, like meats, cheese, and unpasteurized milk. Also, there are other pathways of transmission, like intake of contaminated water, fruits, and vegetables, causing outbreaks of STEC infection [2].

The main virulence factors of STEC associated with HUS are Shiga toxins (Stx). The Stx are AB_5_ toxins, composed of one A subunit and five identical B subunits, which links the toxin to a glycolipid receptor, Gb3 or Gb4, on the surface of the target cells, such as kidney cells, the gastrointestinal tract and central nervous system cells. There are two types of Stx: Stx1 and Stx2, which share approximately 56% of homology in their amino acid sequence. However, Stx2 has been linked epidemiologically to the development of severe diseases, like HUS [3]. Studies on human brain microvascular endothelial cells (HBMC) have shown to be until 1000 times more susceptible to Stx2 than to Stx1 [4], for this reason Stx2 has received greater research attention.

It has been described seven subtypes or genetic variants of Stx2, being Stx2a, Stx2c, and Stx2d the subtypes with the greatest toxicity *in vitro* and the most commonly subtypes identified in STEC strains isolated from patients with clinical manifestations of HUS, unlike the Stx2b, Stx2e, Stx2f and Stx2g subtypes, which have low toxicity and rarely associated with severe diseases in humans, that have been found in animal reservoirs different from cattle like pigs and pigeons [5–6]. The amino acid sequences of the Stx2a, Stx2c, and Stx2d subtypes are closely related, sharing an identity percentage between 97% and 99%, but due to their biological differences and toxicity, they have been classified as different subtypes [7].

The Shiga toxin genes (*stx*) are encoded by temperate bacteriophages inserted in the bacteria chromosome, phages encoding *stx* (*stx* phages) have shown to be variables both morphologically and genetically and play an important role in STEC pathogenesis due they can regulate Stx expression and contribute to the dissemination of the *stx* genes in other bacteria [8]. The *stx* genes are located in the late region of the *stx* phages where genes implicated in the lithic cycle are found. Likewise, late genes are regulated by a transcription anti-terminator protein Q. In absence of the Q protein, the phages cannot carry out their lithic cycle, which implies they will not produce Stx [9]. For the Q protein gene have been identified three allelic variants: *q933*, *q21*, and *qO111,* which are related with differences on the levels of expression of the Stx and virulence of the strains that carrying them [10–12].

In Colombia, the knowledge about STEC is limited and the few studies available have focused on detecting the O157:H7 serotype from different sources and with non-molecular methods [13–14], in addition, no characterization is available of the native strains that have been isolated, either is known their potential virulence. which is why the objective of this work was to characterize biologically and molecularly STEC strains carriers of the stx_2_ gene isolated from bovine cattle in Colombia to provide information on the STEC strains circulating in Colombia, and have an idea of its potential virulence.

## MATERIALS and METHODS

### Bacterial strains

This study was a descriptive cross-sectional, we selected six cattle STEC strains (102, 10610, 600 5052, 615, and N108) positive for the *stx*_*2*_ gene belonging to the collection of strains from the Center of Biomedical Research at Universidad del Quindío. These strains had been characterized previously respect to the presence-absence of STEC virulence genes (*stx*_*2*_, *eae, saa*, and *hly*A) **(Table 1**) (Quiguanas et al 2020, not published). The STEC reference strain EDL933 was used as positive control and the *E. coli* DH5α strain was used as negative control.

**Table 1:** Genetic characteristics and virulence genes of the STEC strains.

| Strain | Virulence genes |  |  | *q* gene alleles |  | *ninG* |
| --- | --- | --- | --- | --- | --- | --- |
|  | Sub Stx2 | *saa* | *hlyA* | *q933* | *qO111* |  |
| **102** | Stx<sub>2d</sub> | + | + | - +<sup>a</sup> | + +<sup>b</sup> | + |
| **10610** | Stx<sub>2a</sub> | + | + | - +<sup>a</sup> | + +<sup>b</sup> | + |
| **5052** | Stx<sub>2c</sub> | + | + | - +<sup>a</sup> | + +<sup>b</sup> | + |
| **600** | Stx<sub>2c</sub> | + | + | - +<sup>a</sup> | - | + |
| **615** | Stx<sub>2d</sub> | + | + | - +<sup>a</sup> | + +<sup>b</sup> | + |
| **N108** | Stx<sub>2c</sub> | + | - | - +<sup>a</sup> | + +<sup>b</sup> | +<sup>c</sup> |

<sup>a</sup> Positive for *primers* QATG and Q3 and negative for *primers* Q-stx-f and 595
<sup>b</sup> Positive for *primers* SF1-F and SF1-R, and SF1-R and 595
<sup>c</sup> Size of amplified 1200 pb

### Induction of the lithic cycle

All the strains were grown individually in Luria-Bertani (LB) broth overnight (ON) at 37 °C, 100 rpm. These cultures were refreshed in LB medium at 37 °C, 180 rpm, when the culture reached the exponential growth phase (Optical Density OD_600_ = 0.2-0.3 nm), it was divided into two subcultures and Mitomycin C (MMC) was added to a fraction at a final concentration of 0.5 µg/ml (inducer of the lithic cycle). The OD_600_ of the cultures both with and without MMC was monitored during five hours to construct growth curves/lysis, growth was measure with a Microplate Spectrophotometer Epoch™ (BioTek). All assays were done at least three times and independently for each strain.

### Stx2 production

To compare the titers of Stx2, the cultures with and without MMC after overnight incubation at 37 °C, 180 rpm were centrifuged at 11.500 rpm, 4 °C during 10 min and the supernatants were evaluated for the presence of Stx2 using an ELISA kit (Ridascreen^®^ Verotoxin, R-Biopharm) according to the manufacturer’s instructions. The ELISA plates were read in the Microplate Spectrophotometer Epoch™ (BioTek) at DO_650_. However, when the DO_650_ values exceeded the limit permitted by the Epoch™ (BioTek), serial dilutions were conducted to obtain a value within the equipment’s reading range. The toxin titers were expressed as the result of the multiplication between the DO_650_ and the reciprocal of the dilution factor; all the assays were carried out at least three times.

### Analysis of the late region of *stx* phages

The upstream region of the *stx*_*2*_ gene was analyzed by PCR in Veriti 96-Well Thermal Cycler (Applied Biosystems™). The proximity of the *ninG* gene with *stx*_*2*_ was evaluated using NinG/526 and 595 *primers* [15], the variability in this gene was evaluated by comparing the sizes of the amplified fragments. The presence of the different alleles of *q* gene (*q933*, *qO111*, and *q21*) was evaluated in all of the strains as well as its closeness with the *stx2* gene. To detect the *q933* allele, the QATG5’ and Q3’ *primers* were used [16], and the closeness with the *stx*_*2*_ gene was evaluated using the Q-stx-f and 595 *primers* [15]. The presence of the *qO111* allele was evaluated with the SF1-F and SF1-R *primers* [12] and its closeness with the *stx*_*2*_ gene was evaluated with the SF1-F and 595 *primers*. The *q21* allele and its closeness with the *stx*_*2*_ gene was evaluated with the Q21 and 595 *primers* [11] (**Fig.1**). All the primers used here are shown in **Table 2**. The conditions for the PCR were the following: 94 °C for 5 min, followed by 30 cycles of 94 °C for 1 min, 60 °C for 1 min, 72 °C for 3 min, ending with 72 °C for 10 min. For the QATG5’ and Q3 *primers* annealing temperature was 53 °C.

**Figure 1.**
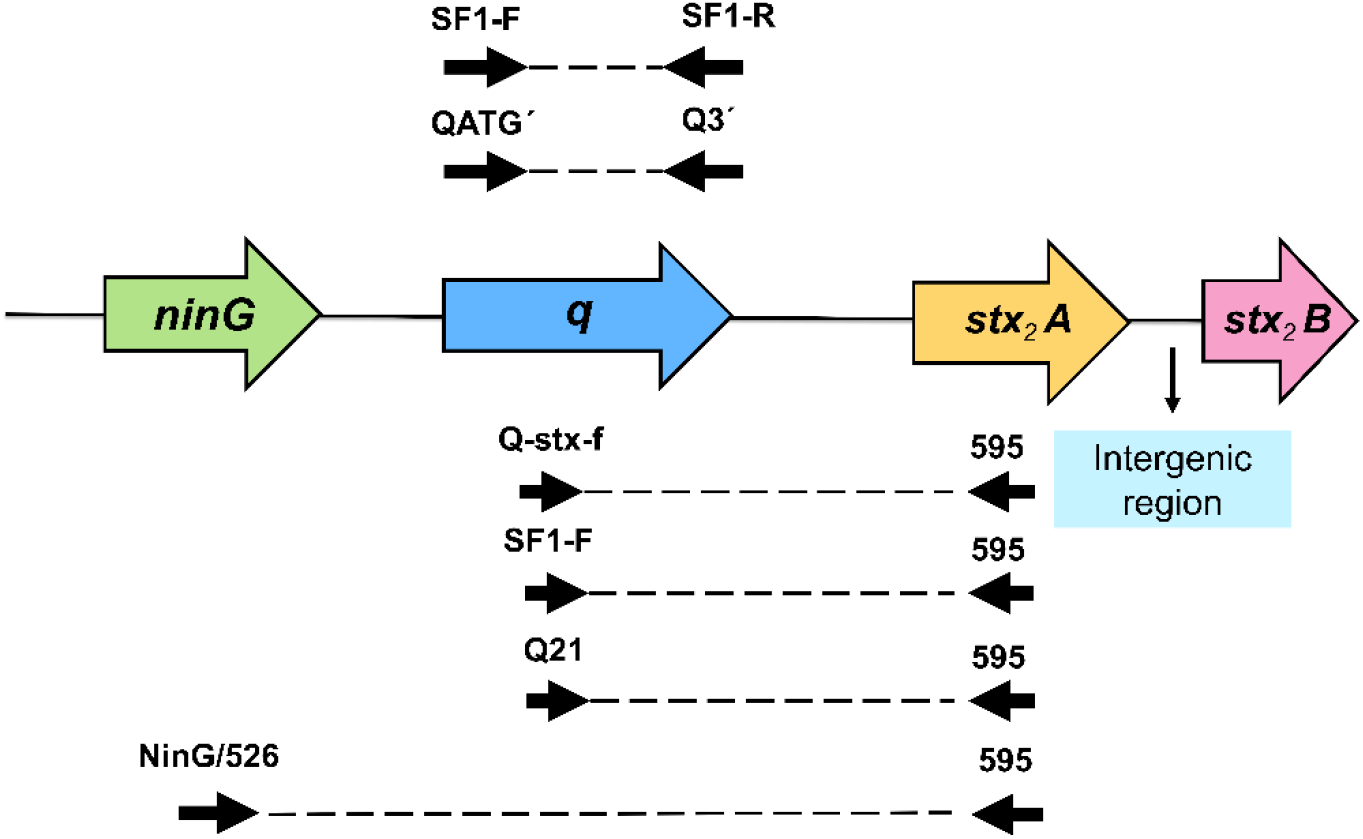
Scheme of the late region of *stx* phages. Colored arrows show the genes evaluated. Dotted lines show the region amplified by the *primers*. Taken and adapted from: Burgan et al [39].

**Table 2.**
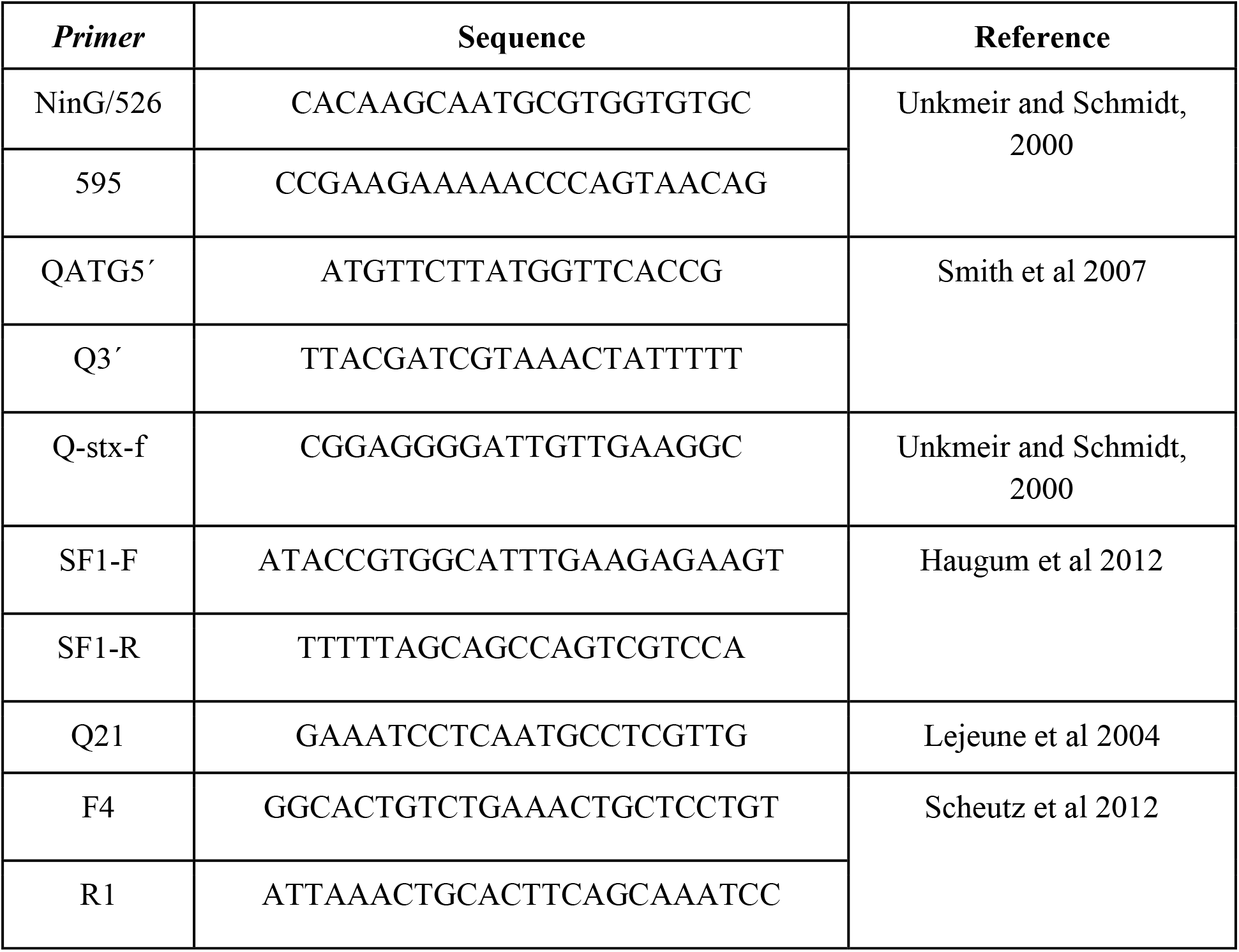
*Primers* used to evaluate the late region and sequencing the *stx_2_* gene of *stx* phages.

### Identification of Stx2 subtypes

The identification of Stx2 subtypes was performed through analysis of sequences with the method proposed by Persson et al [17] and Scheutz et al [7]. A partial sequence of *stx*_*2*_ gene was obtained by using the F4 and R1 sequencing *primers* (**Table 2**), these *primers* amplifying a 491 pb fragment which was translated into 159 amino acids covering 95 residues from the C-terminal region of the A subunit and 64 residues from the N-terminal region of the B subunit, where the sequences of the toxins have greater variability. The PCRs were performed with the Platinum™ Taq DNA Polymerase High Fidelity kit (Invitrogen™) with the following conditions: 95 °C for 15 min, followed by 35 cycles at 94 °C for 50 s, 56 °C for 40 s, and 72 °C for 1 min, and an ending temperature of 72 °C for 3 min. the amplicons obtained with forward and reverse PCR primers were sequenced with the Sanger method in an ABI3500 (Applied Biosystems™). The DNA used for the different PCRs were purified by using the Wizard^®^ Genomics Kit (Promega).

The consensus sequences were obtained with the forward and reverse chromatograms using the Unipro software UGENE RRID: SCR_005579. To identify the subtypes, the intergenic regions between A and B subunits of *stx*_*2*_ sequences of all strains were compared. The nucleotide sequences were translated into amino acids with the ORF established for Stx2 (excluding the intergenic region). A multiple alignment with the amino acid sequences obtained and reference sequences for Stx2 subtypes was carried out by employing the MEGA7 software RRID:SCR_000667 and the Muscle algorithm. Conserved positions described in the literature for each subtype were examined in the multiple alignment; the accession codes for the reference sequences used in the alignment were: Stx2a (GenBank ID: Z37725) Stx2c (GenBank ID: L11079) Stx2d (GenBank ID:DQ059012).

The partial sequences obtained were used to construct a dendrogram by using the UPGMA algorithm (bootstrap of 1000) with various reference sequences of Stx2 subtypes register in the GenBank database: X07865; Z37725; AF524944; AF461173; AY633471; AY633472; EF441599; EF441609; AY443052; AY443057; EF441613; Z50754; DQ344636; FM998856; EF441618; M59432; AB015057; DQ235774; L11079; AY633473; AY443045; AY633467; AY633453; AF291819; EF441604; AY739670; AY739671; AY443044; AY443043; AY443049; AB071845; FM998860; FM177471; EU086525; AF479828; AF479829; AY095209; X61283; DQ235775; AF500190; AF500189; AF500191; AY443047; AY443048; DQ059012; AF329817; AF500192; FM998855; FM998840; EF441605; M21534; AJ313016; M29153; AB472687; AY286000; AB048227; X65949. Although the sequences of the reference strains were complete sequences, in order to perform the analysis, these sequences were cut in the region shown in the alignment (**Fig. 4A**). The sequences obtained in this study were deposited in the GenBank with the following access numbers: MT680394; MT680395; MT680396; MT680397; MT680398; MT680399.

## RESULTS

### Induction of the lithic cycle

Analysis of growth curves showed that all the cultures without MMC had exponential growth with OD_600_ values of ± 1.4, the same behavior was observed with the positive and negative control cultures: EDL933 and DH5α. However, upon inducing the phages with MMC was observed that the 10610 strain had a clear decrease of OD_600_ since the third hour evidencing bacteriolysis by the phage induction. A similar pattern was observed in the 600 strain, but with less decreasing in OD_600_. In the rest of the strains (102, 5052, 615, and N108) MMC caused a decrease in bacterial growth, without evidence of bacteriolysis by phages induction, showing a growth with values of OD_600_ ± 0.8 (**Fig. 2**).

**Figure 2.**
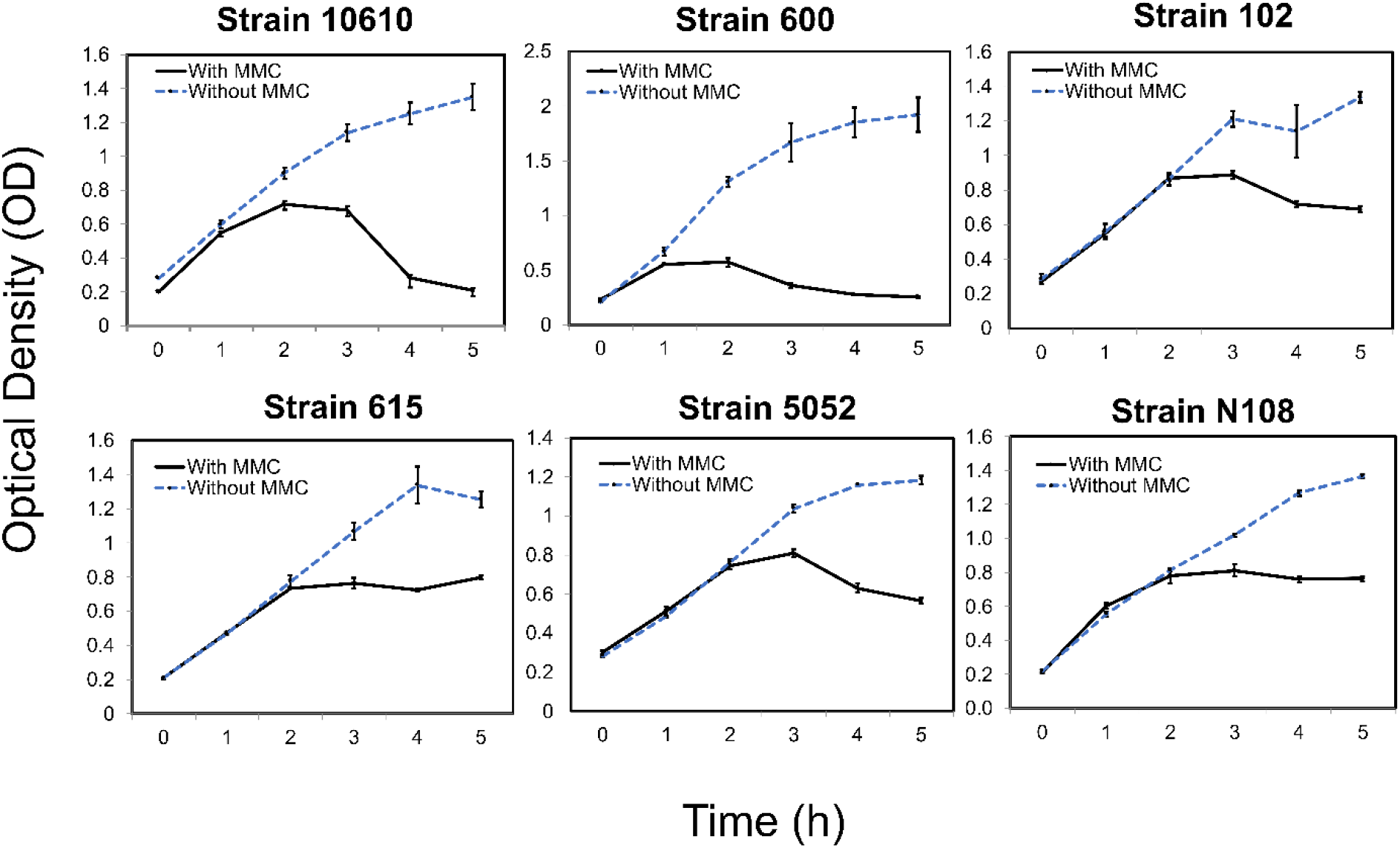
Growth curves/lysis of the strains evaluated in presence/absence of Mitomycin C (MMC).

### Shiga toxin production

All the strains evaluated showed Stx2 production; nevertheless, the titers obtained were different among each other. The 10610 strain showed higher Stx production presenting similar titers to the positive control strain. The 5052, 600, and N108 strains showed lower titers than the control strain, but with moderate Stx production. The 102 and 615 strains produced the lowest titers compared with the other strains (**Fig. 3**). In general, all the strains revealed an increase in the toxin titers when the cultures were induced with MMC. We observed that under non-induction conditions Stx was also detected, but always with low titers, which would demonstrate a basal toxin production in all the strains. The results of absorbance and dilutions carried out are shown in **Table S1.**

**Figure 3.**
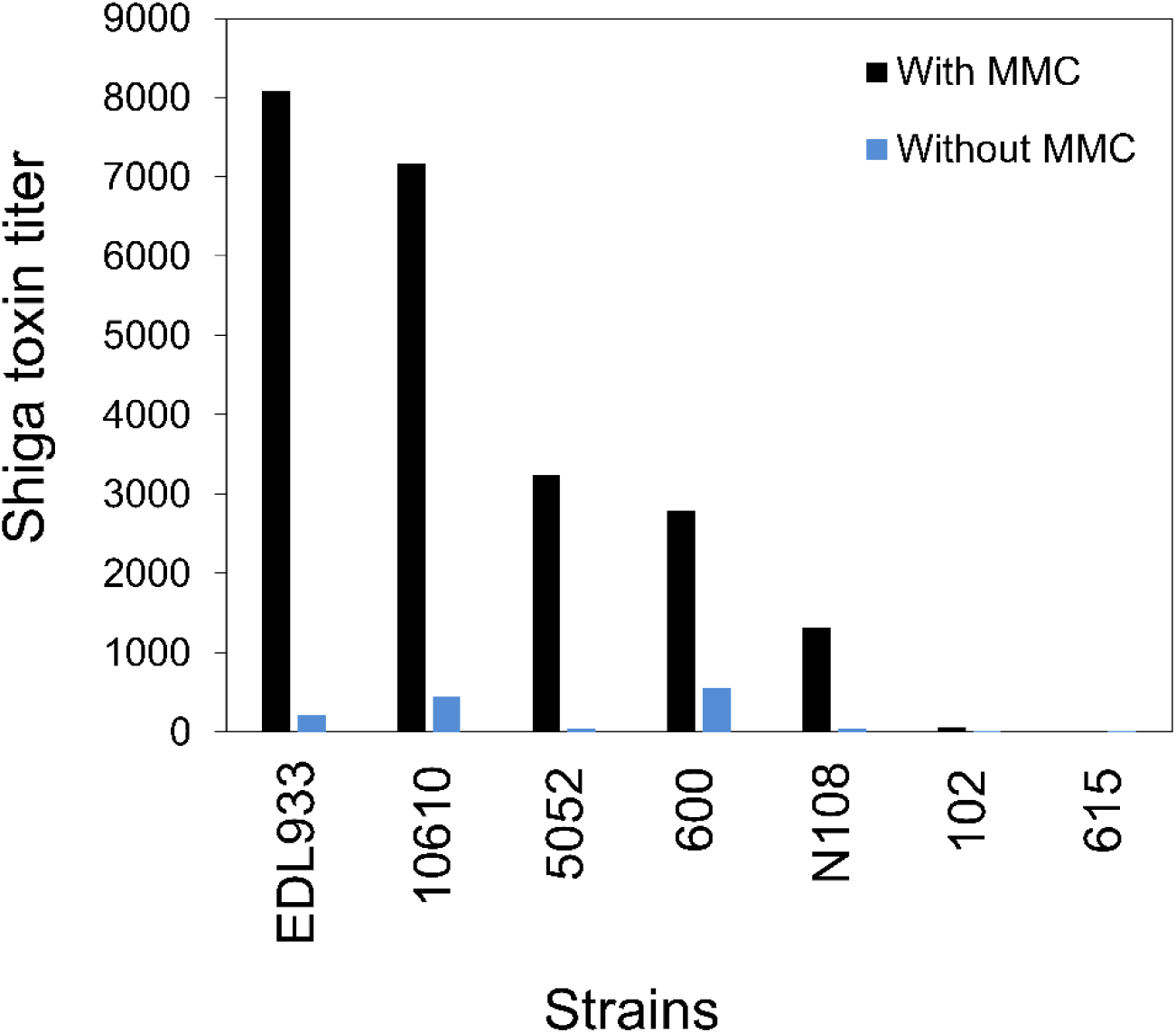
Shiga toxin production of the STEC strains evaluated in presence/ absence of Mitomycin C (MMC).

### Analysis of the late region of *stx* phages

The proximity of the *ninG* gene with the *stx*_*2*_ gene was confirmed in all the strains, where an amplified of 1700 pb fragment was observed, except for the N108 strain that obtained a fragment of 1200 pb suggesting a possible difference in the regulating region of these phages. Regarding the *q* gene alleles, the *q933* allele was detected in all the strains, however, upon evaluating its closeness with the *stx*_*2*_ gene an amplified was not obtained. With respect to the *qO111* allele, its presence and closeness with the *stx*_*2*_ gene were confirmed in five strains. None of the strains studied carried the *q21* allele, the results are shown in **Table 3**.

### Identification of Stx2 subtypes

The chromatograms of some strains showed that in specific positions there were double peaks that corresponded to different nucleotides, in these cases, the IUPAC nucleotide nomenclature was used for resolve these ambiguities. Once the consensus sequence was obtained and translating it into amino acids, it was noted that in most cases the ambiguities in the nucleotides from these positions did not alter the amino acid in the sequence (synonymous mutations); only in two cases (10610 and 600 strains) we observed a codon change in the position 137 and it could be translated into two different amino acids: Alanine or Aspartic acid (**Fig. 4A**).

**Figure 4:**
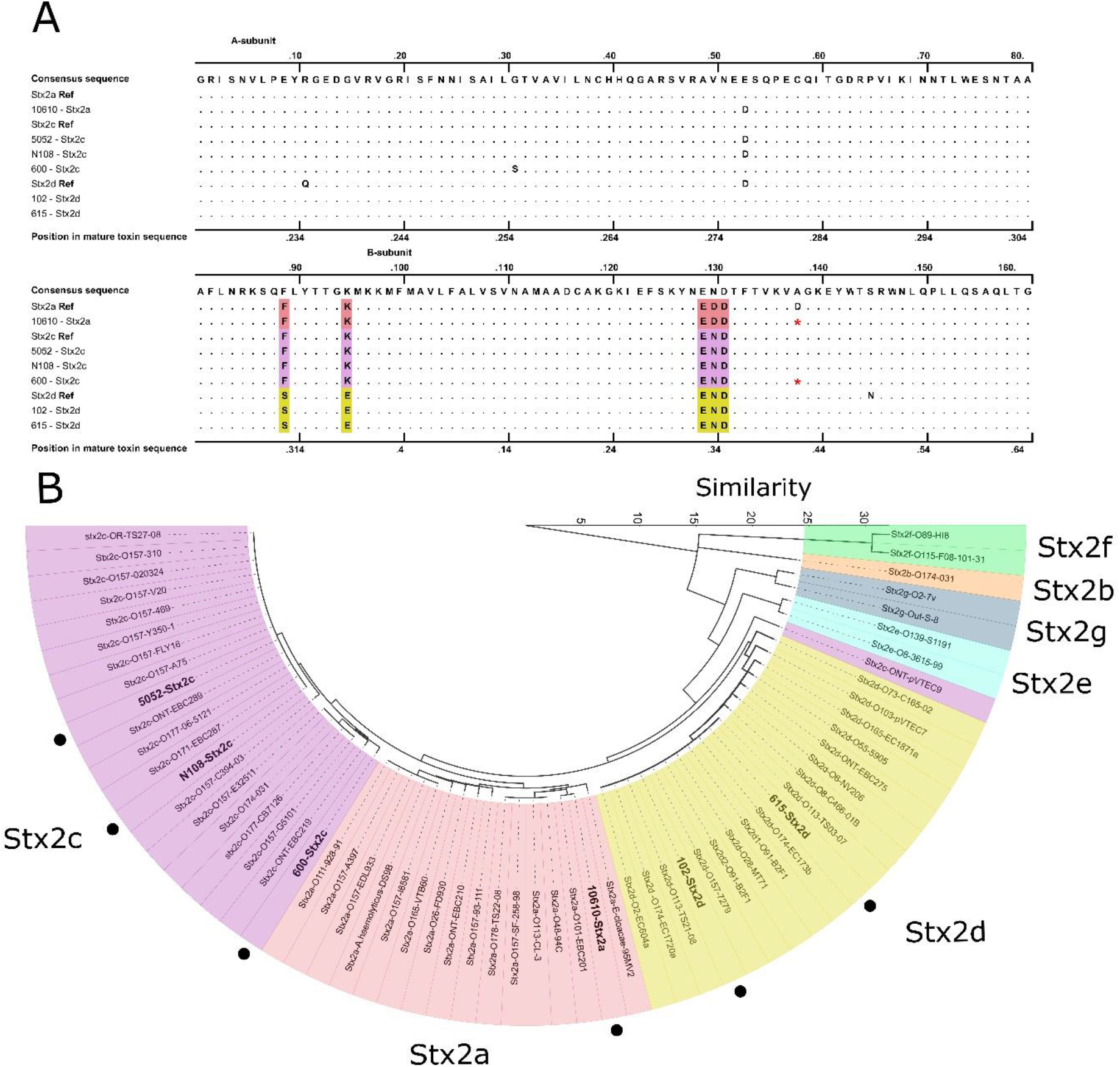
**A.** Multiple alignment with partial reference sequences of Stx2 subtypes and sequences of Stx2 obtained in this study. Colors show conserved sites for each subtype. * In that position both Aspartic acid or Alanine may be present. **B.** The dendrogram shows the groups formed by the reference sequences and the sequences of Stx2 obtained in this study.

Analysis of the intergenic region between the A and B subunit genes of Stx2, showed that all the strains had the same sequence (AGGAGTTAAGT); this sequence has been reported for the Stx2a, Stx2c, and Stx2d subtypes. Analyzing the alignment, we observed that all subtypes sequences were highly conserved and only differ in some positions (**Fig. 4A**) the combination of these positions were used to classify the subtypes, obtaining the following results: Stx2a subtype; one strain, Stx2c subtype; three strains and Stx2d subtype; two strains. Specific changes were observed in amino acid residues in some sequences; however, these are not strictly restricted to a subtype. In the dendrogram both the studied and the reference sequences formed separate groups confirming what was observed in the alignment (**Fig. 4B**),

## DISCUSSION

Presence of inducible phages in the strains studied was evaluated through the analysis of the growth curves/lysis, where it was noted that two of them (10610 and 600) showed bacteriolysis by phage induction with MMC. Likewise, the cultures of these two strains also had the highest Stx titers together with the 5052 strain under induced conditions. Which demonstrates that these strains have phages capable of producing Stx, even at the positive control level. Various studies have shown that the STEC strains with inducible *stx* phages when were treated with MMC increase substantially Stx production, evidencing a direct relationship between induction and Stx production [18–22]. Additionally, Ogura et al [23] have proposed that variability in Stx production among STEC strains may be related with the genetic characteristics of the phages that carry them.

On the contrary, analysis of the growth curves/lysis of the 102 and 615 strains showed a different behavior; both had low response to induction and low Stx titers, even under induced conditions. It has been demonstrated that bacteria can have a high frequency of defective prophages [24–25], which are not capable of carrying out its lithic cycle, limiting its capacity to kill the host bacterial cell; therefore, this would avoid (in the case of *stx* phages) Stx production. Similarly, Johansen et al [26] suggest that the level of Stx production in bacteria carrying apparently defective phages is lower than bacteria carrying inducible phages. In this sense, the low Stx production in the 102 and 615 strains may be explained by the lack of *stx* phage induction observed through the OD kinetics, which could be due to the presence of a defective *stx*_*2*_-phage.

Additionally, in this study we detected Stx production in cultures without MMC, which demonstrates that the strains have a basal expression. In agreement with our results, Gamage et al [27] by analyzing pure supernatants, also reported that all *stx*_2_ strains produced high toxin levels, both in presence and in absence of treatments that induce the phage’s lithic cycle. Probably, the high Stx production under basal conditions is due to a high level of spontaneous induction of the phage that leading to elevated release of Stx. Shimizu et al [28] reported that some *stx* phages have a relatively high spontaneous induction, which results in the presence of Stx2 in the extracellular fraction in absence of any applied induction.

It has been described three alleles encoding for the Q protein (*q933, qO111 and q21)*, which differ in their activity as anti-terminators [10–12]. In this study we detected the *qO111* allele in five strains (102, 10610, 5052, 615, and N108), except the 600 strain. This allele was described recently in Sorbitol-fermenting STEC strains isolated from patients with diarrheic diseases and HUS in several European countries [29–30, 12] which have been related with high progression to HUS [31]. In this study was not possible to demonstrate the closeness of the *q933* allele with the *stx*_*2*_ genes. Perhaps, *q* gene detection in those strains may be due to the presence of defective phages that lack *stx* genes. Regarding the *q21* allele, none strain was found carrying this allele, which has been identified principally in STEC strains with low or null Stx production frequently isolated from beef, animal origin, and the environment in Asian countries, where these are widely distributed [32–34, 10].

Subtyping the STEC strains permitted identifying three Stx2 subtypes: one Stx2a-positive strain, three Stx2c-positive strains and two Stx2d-positive strains, as well as their possible virulence. According to Kawano et al and Fierz et al [35, 5] the STEC strains carriers of these three subtypes have been frequently isolated from patients with HUS, which has led to be strongly related with this disease. The most relevant subtypes identified were: the Stx2a, according to the literature it has demonstrated to be the most potent and active on the endothelial cells [4,6] and the Stx2d, due to an “the activable tail” that causes an increase of up to 25 times its toxicity on Vero cells when it is previously incubated with elastases [36].

Although the three subtypes share a high percentage of identity regarding their amino acid sequence, differences were found in positions 89, 95 on the end of C-terminal of the A subunit and in the END or EDD motif in the B subunit. Fuller et al [6] suggest that the END motif of the B subunit plays an important role in the interaction of the toxin with its receptor, therefore mutations in these positions would cause changes in the affinity of the toxin with his receptor, which could explain the toxicity differences existing among these subtypes. Finally, in the evaluated strains the Stx2b, Stx2e, Stx2g, and Stx2f subtypes were not identified; these subtypes are rarely associated with diseases in humans and contrary to the other subtypes, they have been isolated from sources different from cattle like pigs and pigeons, for that reason they were not detected in this study, due to all the strains were from cattle [37–38].

## CONCLUSION

The results obtained in this study show that the native STEC strains evaluated had differences in the Stx production assays, induction of the phage’s lithic cycle, Stx2 subtypes, and late region of the *stx* phages. Although the sample was limited, it was possible to observe variability among the strains, which could indicate the diversity of the phages carrying the STEC strains in Colombia. Finally, the strains characteristics found in this study are similar to STEC strains that cause diseases in humans reported in the literature. This permit having an idea of the possible virulence of the native STEC, however, more studies are necessary with broader samples that permit determining their virulence in depth.

## KNOWLEDGES

The authors acknowledge to financial support through The Universidad del Quindío through the internal call N°5, project N° 890.

