## Supplementary material for "Biological and molecular characterization of six Shiga toxin-producing *Escherichia coli* (STEC) strains encoding the stx_2_ gene in Colombia": Table S1

[1]Molecular Immunology Group (GYMOL). "Manuel Elkin Patarroyo" Biomedical Research Center. Universidad del Quindío. Armenia. Quindío. Colombia.

**Tabla S1.** Dilutions and absorbances of the ELISA (Ridascreen®Verotoxin R-Biopharm). used to calculate the Shiga toxin titers of the STEC strains evaluated.

| Strains | Shiga toxin production | | | | | |
|---|---|---|---|---|---|---|
| | Dilution | | $OD_{650}$ | | Titer | |
| | With MMC[a] | Without MMC[b] | With MMC[a] | Without MMC[b] | With MMC[a] | Without MMC[b] |
| **102** | 1/100 | 1/10 | 0.63 | 0.76 | 63.9 | 7.68 |
| **10610** | 1/2000 | 1/2000 | 3.58 | 0.22 | 7176.6 | 444.6 |
| **5052** | 1/2000 | 1/100 | 1.61 | 0.37 | 3236 | 37 |
| **600** | 1/4000 | 1/2000 | 0.7 | 0.27 | 2800 | 550.6 |
| **615** | Pura* | Pura* | 3.38 | 0.78 | 3.39 | 0.79 |
| **N108** | 1/500 | 1/100 | 2.63 | 0.29 | 1316.6 | 29.5 |
| **EDL933** | 1/4000 | 1/100 | 2.01 | 2.098 | 8086.6 | 209.8 |

[a.] Supernatan culture with Mitomicyn C (0.5 µg/ml)
[b.] Supernatan culture without Mitomicyn C (0.5 µg/ml)
* Supernatan without dilution
